# Sex differences in the effect of acute fasting on excitatory and inhibitory synapses onto ventral tegmental area dopamine neurons

**DOI:** 10.1101/2020.06.25.172585

**Authors:** Nathan Godfrey, Stephanie L. Borgland

## Abstract

Dopamine neurons in the ventral tegmental area (VTA) are important for energizing goal directed behaviour towards food and are sensitive to changes in metabolic states. Fasting increases the incentive motivation for food, mobilization of energy stores and have sex-dependent effects. However, it is unknown how acute fasting alters excitatory or inhibitory synaptic transmission onto VTA dopamine neurons. An acute 16h overnight fast induced increased food seeking behaviour that was more predominant in male mice. Fasting increased miniature excitatory postsynaptic current (mEPSC) frequency and amplitude in male, but not female mice. This effect was not due to altered release probability as there was no change in the paired pulse ratio, nor was it due to an altered postsynaptic response as there was no change in the AMPAR/NMDA ratio or response to glutamate uncaging. However, this effect was consistent with an increase in the number of release sites. In addition, depolarization-induced-suppression of excitation (DSE), a measure of short-term endocannabinoid-mediated plasticity, was enhanced in female but not male fasted mice. There were no fasting-induced changes at inhibitory synapses onto dopamine neurons of male or female mice. Taken together, these results demonstrate that fasting influences excitatory synapses differentially in male and female mice, but preserves inhibitory synapses onto dopamine neurons, indicating that the mesolimbic circuit of male and female mice respond differently to acute energy deprivation.

**Key Points:**

1. Fasting can increase motivation for food and can energize reward-seeking.
2. Ventral tegmental area (VTA) dopamine neurons respond to motivationally relevant information and fasting can influence mesolimbic dopamine concentration.
3. An acute overnight fast differentially alters food approach behaviours and excitatory synaptic transmission onto VTA dopamine neurons of male or female mice.
4. While inhibitory synapses onto VTA dopamine neurons are not altered by fasting in male or female mice, male mice had strengthened excitatory synapses whereas female mice had increased endocannabinoid-mediated short term plasticity at excitatory synapses.
5. These results help us understand how fasting differentially influences excitatory synaptic transmission onto dopamine neurons and may inform different strategies for fasting-induced food seeking by male and female mice.

## Introduction

Increased obesity rates have been paralleled with increased dieting amongst the general population (Julia *et al*., 2014). While achieving modest body weight reduction can improve cardiovascular and glycemic outcomes (Wing *et al*., 2011), weight loss is rarely maintained. Hypotheses have been put forward to understand why permanent weight loss is seldom achieved. First, obesogenic diets can induce long lasting plasticity in neural circuits that motivate food intake and drive feeding behaviour (La Fleur *et al*., 2007; Liu *et al*., 2016). Secondly, food restriction and weight loss can induce metabolic and neural changes that enhance reward-seeking and food intake (Fulton, 2010; Sharma *et al*., 2013; Matikainen-Ankney *et al*., 2020). Ventral tegmental area (VTA) dopamine neurons encode motivationally salient information, and dopamine released in the nucleus accumbens enhances motivation for food (Salamone & Correa, 2012). Increased synaptic strength of excitatory inputs in the VTA underlies learning of reward-associated cues (Stuber *et al*., 2008). Acute exposure to sweetened high fat food strengthens excitatory synapses onto VTA dopamine neurons and drives further food consumption (Liu *et al*., 2016). However, it is unclear how food restriction and weight loss influence synaptic plasticity in the VTA. Furthermore, a greater understanding of mechanisms underlying food restriction or fasting may inform why weight loss is rarely maintained and perhaps shed light on how eating disorders may develop.

Both chronic food restriction (Reilly, 1999; Carr, 2002) and acute fasting (Jewett *et al*., 1995) increase motivation for food. Numerous studies have shown that chronically food restricted mice and rats have greater dopamine release in the nucleus accumbens (Heffner *et al*., 1980; Wilson *et al*., 1995; Cadoni *et al*., 2003). In addition, chronically food restricted male mice have increased burst firing of dopamine neurons in the substantia nigra (Branch *et al*., 2013). This is associated with an increased excitatory synaptic strength of dopamine neurons due to an increase in the AMPAR/NMDAR ratio, as well as increased sensitivity of dopamine D2 autoreceptors to activation dependent desensitization (Branch *et al*., 2013). Conversely, overnight fasting does not affect burst firing or the AMPAR/NMDAR ratio of substantia nigra dopamine neurons from male mice (Branch *et al*., 2013). Acute fasting (24-36h) decreases activity of dopamine transporter and decreases dopamine reuptake in the rat striatum (Paterson *et al*., 1998). In the VTA, a 24 h fast increases somatodendritic dopamine D2 receptor-mediated inhibitory post synaptic current (IPSC) responses (Roseberry, 2015). Thus, while fasting can influence dopamine signaling, there is a lack of understanding how acute fasting can influence excitatory and inhibitory synaptic transmission within the VTA. Furthermore, while there are reported sex differences in the physiological response to fasting in humans (Johnstone *et al*., 2002) and mice (Jensen *et al*., 2013; Freire *et al*., 2020), it is not known how fasting influences synaptic transmission in the VTA differently between male and female rodents. Therefore, we hypothesized that acute overnight fasting influences excitatory and inhibitory synaptic transmission onto VTA dopamine neurons differently in male and female mice.

## Methods

### Ethical Approval

All experiments and procedures were in accordance with the ethical guidelines established by the Canadian Council for Animal Care and were approved by the University of Calgary Animal Care Committee (protocol number AC17-0004) and are consistent with the journal’s policies for the ethical treatment of animals (Grundy, 2015).

### Animals

Mice 2-4 months old were housed in groups of 2-5 in same sex cages and were maintained on a 12 h light-dark schedule (lights on at 8 AM MST, zeitgeber time (ZT0)) and were given chow and water *ad libitum*. Mice were fed chow (5062 from Pico-Vac), which is composed of (% of total kCal) 23.189 protein, 21.635 fat (ether extract), and 55.176 carbohydrate. The total energy density of this diet was 4.60 kCal/g. Experiments were performed during the animal’s light cycle. Male and female DAT^cre^Td-Tomato mice, used to identify dopamine neurons, were generated by crossing DAT-Cre (B6.SJL-Slc6a3tm1.1(cre)Bkmn/J mouse line) with Rosa-td Tomato mice (B6.Cg-Gt(ROSA)26Sortm9(CAG-tdTomato)Hze/J (Ai9)). All mice were bred locally in the Clara Christie Centre for Mouse Genomics. Mice were initially weighed and then isolated 10 hours into their light phase (ZT10), and single housed overnight for a total of 16 hours. During this time, food was removed from the cages of the fasted group and chow was given *ad libitum* to the control group. Water was given *ad libitum* to both groups. Water intake and body weight were recorded for fasted and control groups, and food consumption per body weight was recorded in the control group after the 16h fast (ZT2). In a separate cohort of mice, at ZT1, glucose, ketone, and serum corticosterone were measured from tail vein blood. Electrophysiology or behavioural experiments were performed on mice at ZT2.

### Ketone, Glucose and CORT measurements

Glucose measurement was performed using a glucose meter (Accu-check Aviva, Roche) on a drop of tail vein blood. Ketone measurement was performed similarly on a drop of tail vein blood, using a ketone blood monitor with keto testing strips (Freestyle Precision Neo, Abbott Laboratories Inc.). Blood serum was separated after centrifuging blood samples, obtained from the tail using Microvette collection tubes (Microvette CB 300 uL, Clotting Activator/Serum tubes, Sarstedt, Inc.) at 16,100 g, 15 min at 4 °C and serum corticosterone (CORT) was measured with and according to manufacturer’s instructions (DetectX Corticosterone Enzyme Immunoassay Kit, Arbor Assays).

### Food-seeking behaviour

Mice were pre-exposed to a ∼5 g high-fat diet pellet 48 hours prior to the behavioural test to reduce food neophobia. This high-fat food (D12492 from Research Diets) is composed of (% of total calories) 20 protein, 60 fat, and 20 carbohydrate. The total energy density of this diet was 5.21 kcal/g. Following isolation, fasted or control mice were individually placed in a light-dark box apparatus for 10 minutes. The apparatus was an acrylic box (40 cm length x 40 cm width x 12 cm height) divided equally into light (60 lumens) and dark (5 lumens) chambers with an opening (3.5 cm x 3.4 cm) for the mouse to move from zone to zone. A pellet of high-fat food (Research Diets, D12492) was placed at the centre of the light side, and the area around the food (9 cm x 9 cm) was defined as the food zone (Liu *et al*., 2016). Mice were placed in the light chamber facing the opening into the dark chamber. Movement of the mouse was recorded and analyzed using EthoVision XT software (Noldus) as well as MatLab (code:https://github.com/borglandlab/Godfrey-and-Borgland-2020.git). The apparatus was cleaned between each subject. Mice were then given access to the high fat diet for an hour following the light dark box test, and consumption was measured.

### Electrophysiology

All electrophysiological recordings were performed in slice preparations from adult (2-4 months old) DAT^cre^;Td-Tomato mice. In these mice, a red fluorescent marker is expressed exclusively in dopamine neurons, permitting immediate identification of each recorded neuron in the VTA. In all electrophysiology experiments, we recorded fluorescently-identified dopamine neurons from the lateral VTA located medial to the medial terminal nucleus of the accessory optic tract (MT). Mice were deeply anaesthetized with isoflurane and transcardially perfused with an ice-cold NMDG solution containing (in mM): 93 NMDG, 2.5 KCl, 1.2 NaH_2_PO_4_.2H_2_O, 30 NaHCO_3_, 20 HEPES, 25 D-Glucose, 5 Sodium Ascorbate, 3 Sodium Pyruvate, 2 Thiourea, 10 MgSO_4_.7H_2_0, and 0.5 CaCl_2_.2H_2_0 and saturated with 95% O_2_/5% CO_2_. Mice were then decapitated, and brains were extracted. Horizontal midbrain sections (250 µm) containing the VTA were cut on a vibratome (Leica, Nussloch, Germany). Slices were then incubated in NMDG solution (32°C) and saturated with 95% O_2_/5% CO_2_ for 10 minutes. Following this, the slices were transferred to aCSF containing (in mM): 126 NaCl, 1.6 KCl, 1.1 NaH_2_PO_4_, 1.4 MgCl_2_, 2.4 CaCl_2_, 26 NaHCO_3_, 11 glucose (32°C) and saturated with 95% O_2_/5% CO_2_ and incubated for a minimum of 45 min before being transferred to a recording chamber and superfused (2 mL/min) with aCSF (32-34°C) and saturated with 95% O_2_/5% CO_2_. Cells were visualized on an upright microscope using “Dodt-type” gradient contrast infrared optics and whole-cell recordings were made using a MultiClamp 700B amplifier. The recorded signal was collected at a sampling rate of 20 KHz. Except for recordings of miniature post-synaptic currents (mPSC), a 2 KHz Bessel filter was applied to the data during collection using the MultiClamp 700B (Molecular Devices, CA). For mPSC recordings, a 10 KHz Bessel filter was applied to the data during collection and the data was post hoc filtered at 2 KHz after the recordings using Clampfit 10.3 (Molecular Devices, CA). For evoked currents, a bipolar stimulating electrode was placed 100 to 300 µm rostral to the cell being recorded and stimulated at 0.1 Hz. The liquid junction potential was calculated using a tool on Clampex, but not corrected for.

Recording electrodes (2-5 MΩ) were filled with CeMeSO_3_ internal solution containing (in mM): 117 CeMeSO_3_, 2.8 NaCl, 20 HEPES, 0.4 EGTA, 5 TEA, 5 MgATP, 0.5 NaGTP. The liquid junction potential was -10 mV. For miniature excitatory post-synaptic currents (mEPSC), cells were voltage clamped at the reverse potential of inhibitory currents, experimentally determined to be -65 mV, and the current was recorded for 5 minutes. For miniature inhibitory post-synaptic currents (mIPSC), cells were voltage clamped at the reverse potential of excitatory currents, experimentally determined to be +9 mV, and the current was recorded for 5 minutes. mEPSCs and mIPSCs were identified and measured using Mini Analysis 60 (Synaptosoft, Decateur GA) and the following parameters: mEPSC: amplitude > 12 pA, decay time < 3 ms, rise time < 1.75 ms; mIPSC: amplitude > 15 pA, decay time < 10 ms, rise time < 4 ms.

Recording electrodes (2-5 MΩ) were filled with CeMeSO_3_ internal solution and the evoked currents were recorded in the presence of picrotoxin (100 µM). The α-amino-3-hydroxy-5-methyl-4-isoxazolepropionic acid receptor (AMPAR) to N-methyl-D-aspartate receptor (NMDAR) ratio was recorded while the cells were voltage clamped at +40 mV, and a single evoked stimulus was delivered. Using Matlab (code: https://github.com/borglandlab/Godfrey-and-Borgland-2020.git) the average of 12 current amplitudes, over 2 min, was calculated before and after the application of the NMDAR receptor antagonist, AP5 (50 µM). NMDAR current amplitude was calculated by subtracting the current amplitude after AP5 (AMPAR only) from the current amplitude without AP5. The AMPAR EPSC amplitude was divided by the NMDAR EPSC amplitude to calculate the AMPAR/NMDAR ratio (Borgland *et al*., 2006). To elicit glutamatergic current via AMPARs voltage clamped at -70 mV, Rubi-Glutamate (30 µM) was washed on to the slice. Glutamate uncaging was triggered by a 15 ms, 3.5 mW, pulse of 470 nm, provided by an LED light source through a 40X/0.80 water immersible Olympus microscope objective every minute (0.02 Hz) (Fino *et al*., 2009). Recorded cells were positioned so that the soma was at the centre of the objective. Averaged current amplitudes were an average of 5 pulses over 5 minutes.

To examine short term plasticity, EPSCs recorded at -70 mV were electrically evoked using 40 pulses at 100 Hz. Using Matlab, the amplitudes of the first 2 stimulations were averaged and the cumulative probability of amplitude was plotted, and a line was fitted to the final 14 pulses. The readily releasable pool (RRP) was calculated as the y-intersect of this best fitting line. The probability of release (p) was calculated by dividing the first peak (EPSC_0_) by the RRP. The PPR was calculated by dividing the second peak (EPSC_1_) by EPSC_0_ (Thanawala & Regehr, 2013, 2016).

### Depolarization-induced suppression of excitation

To evoke depolarization-induced suppression of excitation (DSE), neurons were voltage clamped at -70 mV in the presence of picrotoxin (100 µM), and a paired stimulus was delivered with an interstimulus interval (ISI) of 50 ms. Prior to each 10 s depolarization step to +40 mV, evoked EPSCs were recorded for 1 min, and the resulting 6 EPSCs from this time were averaged to create a baseline (Melis *et al*., 2004). The magnitude of DSE was measured as the percentage of the amplitude of the EPSC immediately after depolarization (acquired between 5 and 15 s after the end of the pulse) relative to the baseline before depolarization. The depolarization step occurred a minimum of 3 times, and an averaged DSE was calculated from these steps. To confirm DSE was mediated by endocannabinoids, the protocol was repeated in the presence of the CB1R antagonist, AM251 (2 µM). The paired-pulse ratio was calculated by dividing the second pulse by the first pulse during the baseline.

### Application of WIN 55,212

A maximal concentration of CB1 receptor agonist, WIN 55,212 (10 µM) was applied to VTA slices following a 10 min baseline. A paired stimulus was delivered with an interstimulus interval (ISI) of 50 ms. The 30 EPSCs from the 5 min prior to application of WIN 55,212 were averaged to create a baseline. EPSCs were recorded for 20 min following WIN 55,212 application and the final 30 EPSCS from the last 5 minutes of the recording were averaged and compared to baseline. The paired-pulse ratio was calculated by dividing the second pulse by the first pulse during both the 5 min baseline and final 5 min of the recording.

### Depolarization-induced suppression of inhibition (DSI)

Recording electrodes (2-5 MΩ) were filled with a KCl internal solution containing (in mM): 144 KCl, 10 HEPES, 2.6 BAPTA tetrapotassium salt, 1 CaCl_2_, 2.5 Mg_2_ATP, 0.25 Na_2_GTP, 3 QX 314 – Cl. The liquid junction potential was -4 mV. Neurons were voltage clamped at -70 mV and a paired stimulus (ISI 50 ms) was delivered in the presence of DNQX (10 µM) and AP5 (50 µM). Prior to each 5 s depolarization step to +40 mV, evoked IPSCs were recorded for 1 min, and the resulting 6 IPSCs from this time were averaged to create a baseline (Melis *et al*., 2013). The magnitude of DSI was measured as the percentage of the amplitude of the IPSC immediately after depolarization (acquired between 5 and 15 s after the end of the pulse) relative to that of the baseline before depolarization. The depolarization step occurred a minimum of 3 times, and an averaged DSI was calculated from these steps. To confirm DSI was mediated by endocannabinoids, the protocol was repeated in the presence of the CB1R antagonist, AM251 (2 µM). The paired-pulse ratio was calculated by dividing the second pulse by the first pulse during the baseline.

### Data Analysis

All values are expressed as mean ± SD and assessed for normality using a Shapiro-Wilk test. In figures 1 and 2 symbols represent individual mice, while in figures 3 to 9 symbols represent individual cells. Statistical significance was assessed by using two-tailed unpaired Student’s t test for 2 comparisons. A two-way ANOVA followed by a Tukey’s multiple comparisons was used for multiple group comparisons. For time course experiments, a repeated measures two-way ANOVA followed by Sidak’s multiple comparisons test were used. In all electrophysiology experiments, sample size is expressed as N/n, where N refers to the number of cells recorded from n animals. Asterisks were used to express statistical significance in figures: *P<0.05, **P<0.01, ***P<0.001, and ****P<0.0001. GraphPad Prism 8.3 was used to perform statistical analysis. Unless otherwise stated, figures were generated using GraphPad Prism 8.3 and Adobe Illustrator CS4 software (Adobe Systems Incorporated).

**Figure 1.**
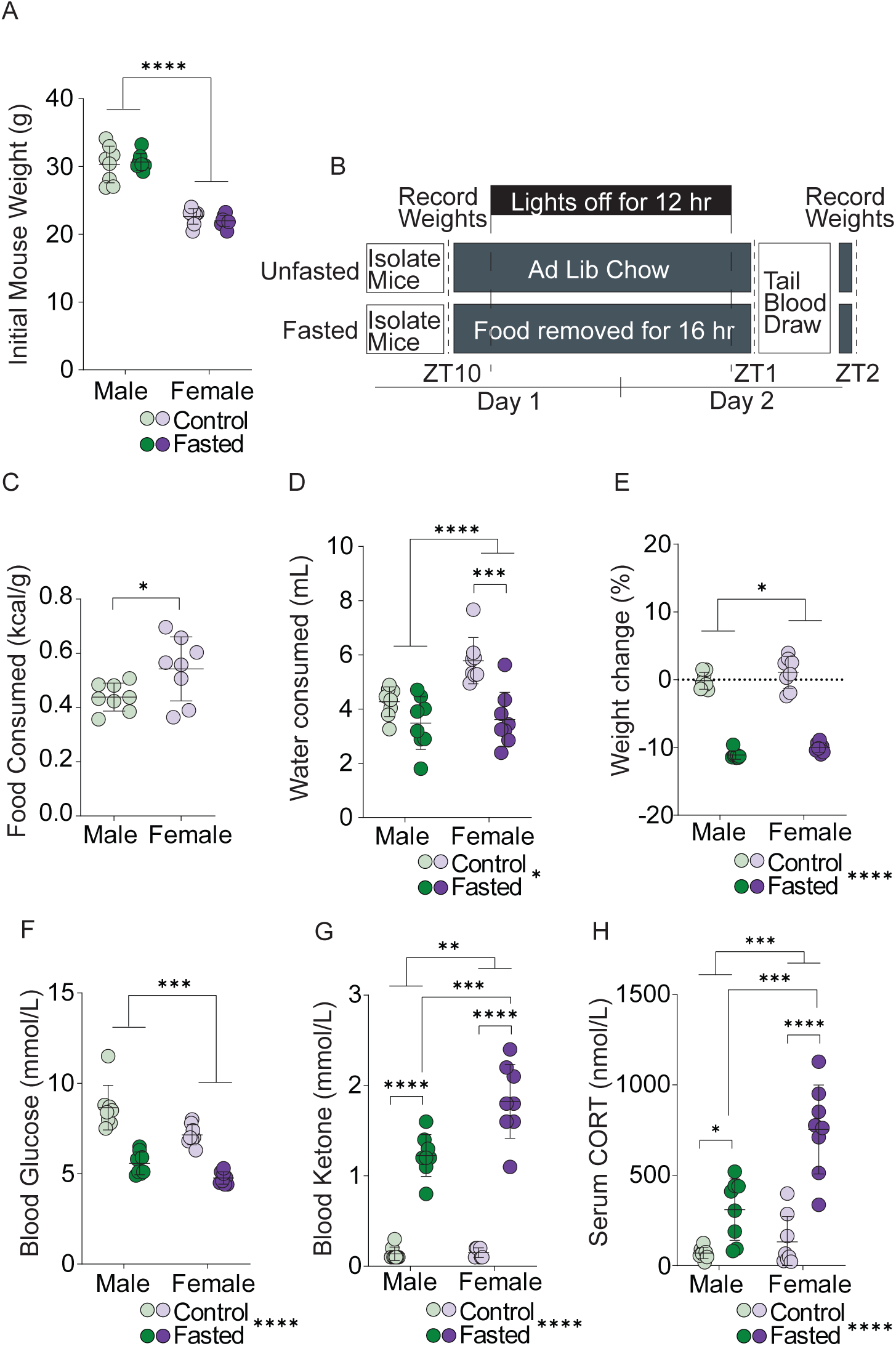
Body weight and metabolic measures after acute fasting in male and female mice. A) Initial mouse weight (g) prior to the 16 h overnight isolation or fast. Male mice weigh more than female mice. B) Schematic showing time course of fast and when blood draw and weights were recorded. C) Chow consumed (kcal/body weight (g)) by control male and female mice during the 16h overnight period of isolation. Female mice consume more food per body weight than male mice. D) Water consumed by male and female mice during the 16h overnight period. Female mice reduce water intake during the acute fast. E) Body weight change between before and after the 16h overnight period. Male and female fasted mice have reduced bodyweight after fasting. F) Blood glucose concentration (mmol/L) measured after the 16h overnight period. Male and female fasted mice have decreased blood glucose. G) Blood ketone concentration (mmol/L) measured after the 16h overnight period. Male and female fasted mice have increased blood ketones. H) Serum CORT concentration (mmol/L) measured after the 16h overnight period. Male and female fasted mice have increased serum CORT, this effect was greater in female mice.

**Figure 2.**
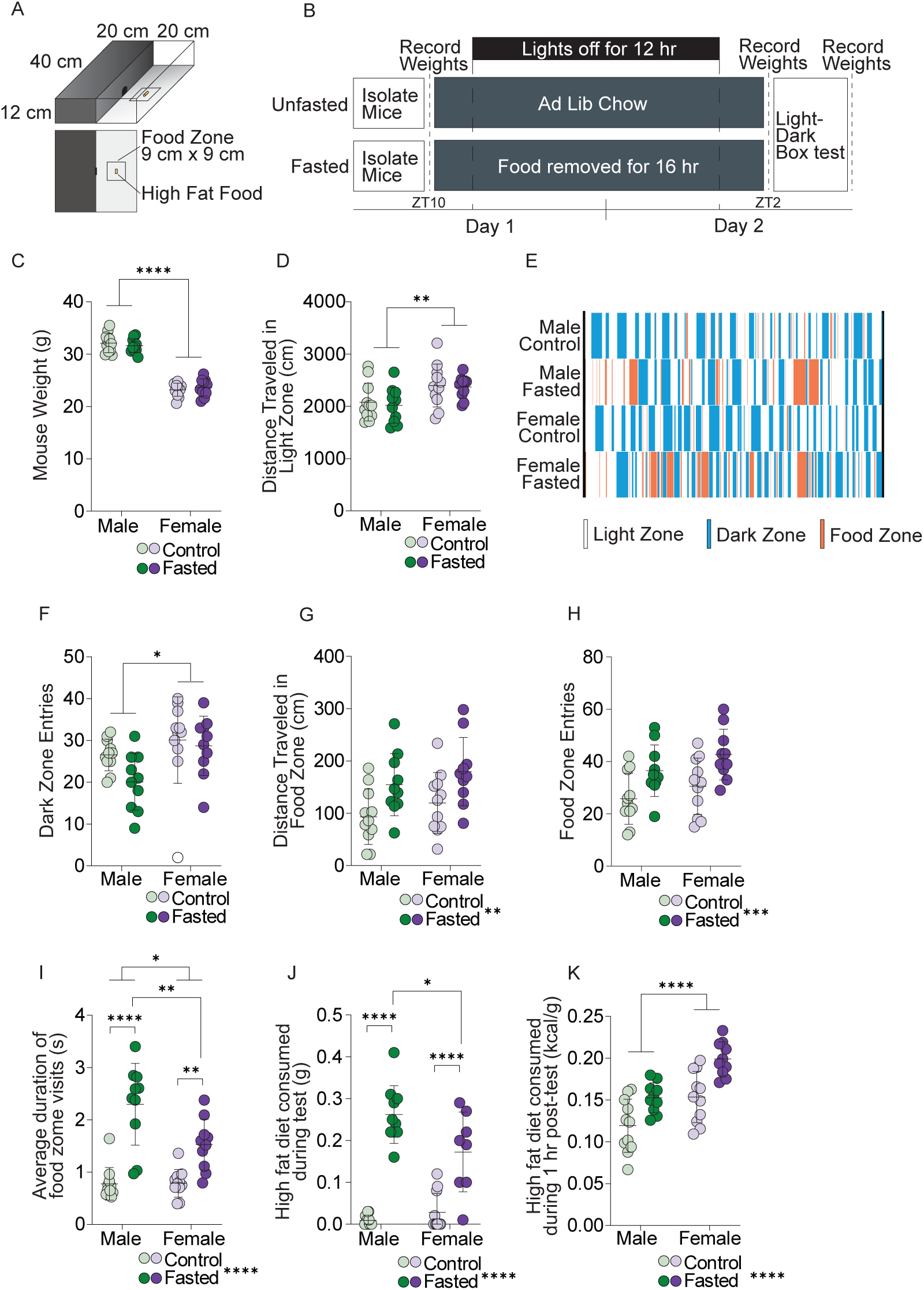
Sex differences in food approach behaviour and food consumption following fasting. A) Schematic of the light-dark box apparatus with the food zone. B) Schematic showing time course of fast and when behavioural performance was recorded. C) Mouse weight (g) prior to the behavioural task. Male mice weigh more than female mice. D) Distance travelled in the light zone during the 10 min test after fasting. Female control or fasted mice had increased distance traveled compared to males. E) Plot illustrating behavior in the light zone (shaded bars), dark zone (filled bars) or food zone (open bars) during the 10 min light-dark box test of individual representative mice. F) Dark zone entries during the 10 min test after fasting. Females had increased dark zone entries compared to males. G) Distance travelled in the food zone during the 10 min test after fasting. Fasting increased distance traveled in the food zone. H) Food zone entries during the 10 min test after fasting. Fasting increased food zone entries. I) Average duration of food zone visits during the 10 min test after fasting. Fasting increased the average duration in the food zone in male and female mice. Male fasted mice had longer food zone visits than female fasted mice. J) High fat diet consumed (g) during the 10 min test after fasting. Fasting increased food consumed during the test. Male fasted mice consumed more than female fasted mice. K) High fat diet consumed (kcal/body weight (g)) in 1 hr following the light-dark box test. Female mice consume more kcal/g than male mice.

**Figure 3.**
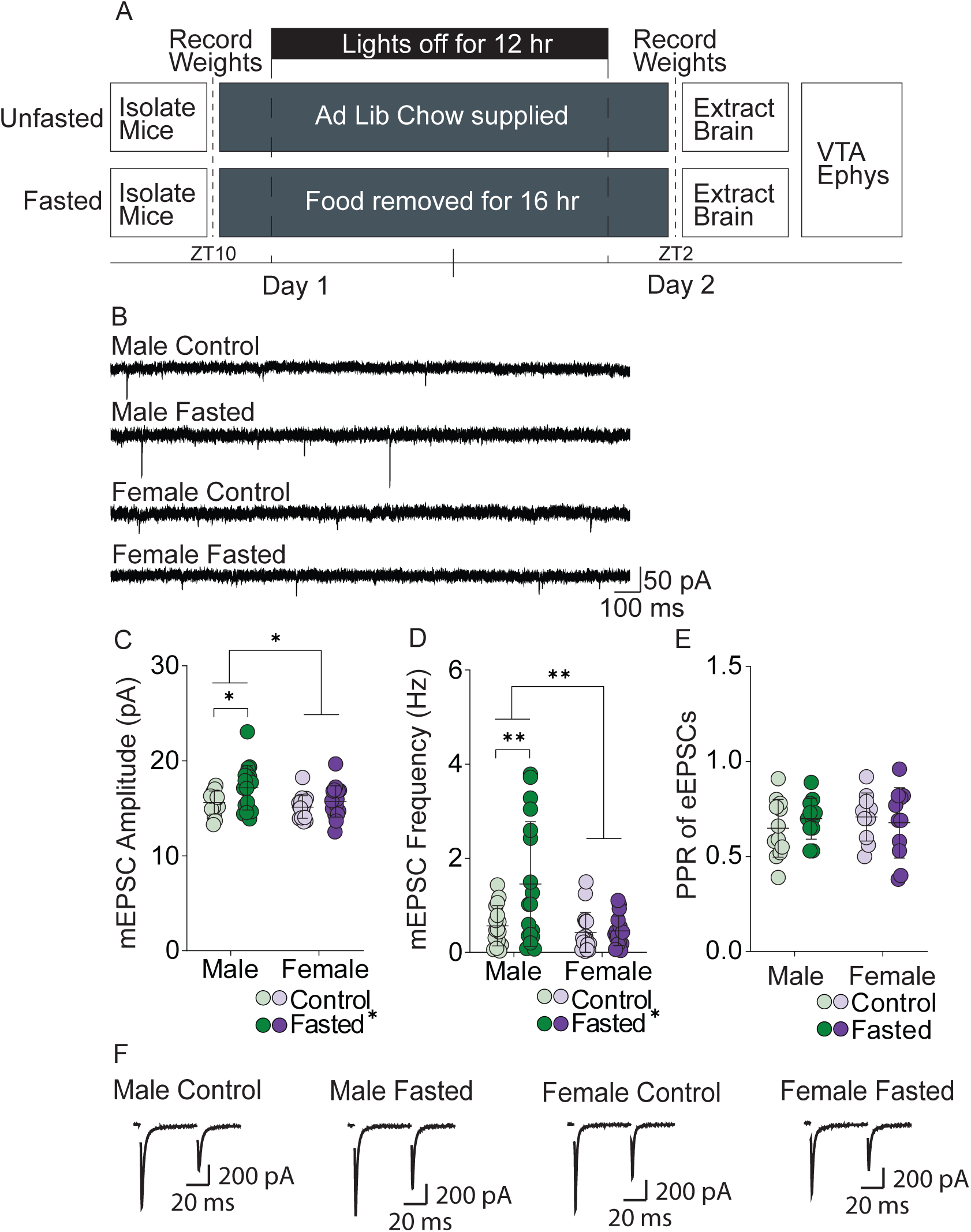
Sex differences in quantal release properties at excitatory synapses onto VTA dopamine neurons of male and female mice following fasting. A) Schematic showing time course of fast and when brain slices were extracted for electrophysiology experiments. B) Representative examples of mEPSC recordings from male and female fasted and control mice. C) mEPSC amplitude onto dopamine neurons of male and female fasted and control mice. Fasting increased mEPSC amplitude in male mice, but not female mice. D) mEPSC frequency onto dopamine neurons of male and female fasted and control mice. Fasting increased mEPSC frequency in male mice, but not female mice. E) Paired-pulse ratio of evoked EPSCs was not different between fasted or control male and female mice. Fasting did not alter PPR in male or female mice. F) Representative example recordings of the paired-pulse evoked EPSC currents.

## Results

### Changes in body weight and concentrations of blood glucose, blood ketones, and serum CORT following fasting

To characterize the physiological effects of a 16 h overnight fast on DAT^cre^Td-Tomato mice, changes in weight, food and water consumption, blood glucose concentration, blood ketone concentration, and serum CORT concentration were measured (Figure 1A). All mice were weighed prior to the 16 h overnight isolation/fast, and a main effect of sex was observed but no difference between fasted and control groups (sex effect: F(1,28) = 197.6, P < 0.0001; male fasted: 30.7 ±1.2 g, n = 8; female fasted: 23.0 ±0.9 g, n = 8; male control: 30.3 ±2.7 g, n = 8; female control: 22. ± 1.1 g, n = 8; Figure 1A). Despite differences in initial body weight, control mice consumed a similar quantity of food during their 16 h overnight isolation, resulting in female mice consuming more kcal/body weight (g) than male control mice (t(14) = 2.283, P = 0.0386; male control: 0.44 ± 0.051 kcal/g, n = 8; female control: 0.54 ±0.12 kcal/g, n = 8; Figure 1C). Fasting decreased water consumption in male (control: 4.3 ±0.5 mL, n = 8; fasted: 3.5 ±1.0 mL, n = 8) and female (control: 5.8 ±0.9 mL, n = 8; fasted: 3.6 ±1.0 mL, n = 8,) mice (sex x fasting interaction: F(1,28) = 5.140, P = 0.0313; fasting effect: F(1,28) = 7.302, P = 0.0116). Furthermore there was a sex difference in water consumption (sex effect: F(1,28) = 23.55, P < 0.0001; Figure 1D). A Tukey’s multiple comparison test indicated a significant difference in water consumption between female (P<0.0001) but not male (P = 0.2818) control and fasted mice. Male (control: -0.2 ± 1.2 %, n = 8, fasted = -11.1 ±0.6 %, n = 8) and female (control = +1.1 ± 2.3 %, n = 8; fasted = -10.0 ± 0.7 %, n = 8) mice lost weight during fasting (fasting effect: F(1,28) = 495.2, P < 0.0001, sex effect: F(1,28) = 5.695, P = 0.0240, sex x fasting interaction: F(1,28) = 0.0257, P = 0.8738; Figure 1E).

Following fasting, there was a main effect of fasting on blood glucose concentration in male and female mice (F(1,28) = 101.6, P < 0.0001, male control: 8.6 ±1.2 mmol/L, n = 8; female control: 7.1 ±0.6 mmol/L, n = 8, male fasted: 5.6 ± 0.6 mmol/L, n = 8; female fasted: 4.8 ±0.3 mmol/L, n = 8), and a main effect of sex (F(1,28) = 18.20, P = 0.0002; Figure 1F). There was no sex x fasting interaction (F(1,28) = 1.609, P = 0.2151). There was a main effect of fasting on blood ketone concentrations (F(1,28) = 265.6, P < 0.0001, male control: 0.1 ±0.07 mmol/L, n = 8; female control: 0.15 ±0.05 mmol/L, n = 8, male fasted: 1.2 ±0.2 mmol/L, n = 8; female fasted: 1.8 ±0.4 mmol/L, n = 8). Furthermore, there was a main effect of sex on blood ketones (F(1,28) = 13.06, P = 0.0012; fasting x sex interaction: F(1,28) = 12.01, P = 0.0017). A Tukey’s multiple comparison test revealed a significant difference between ketone levels between male and female fasted mice (P = 0.0002), but not control mice (P = 0.9996, Figure 1G). Finally, there was a significant sex x fasting interaction on serum CORT concentrations (F(1,28) = 10.60, P = 0.0030, main effect of fasting: (F(1,28) = 53.96, P < 0.0001), main effect of sex (F(1,28) = 18.46, P = 0.0002). A Tukey’s multiple comparison’s test reveals a significant difference in CORT levels between control and fasted male (P = 0.0348, control: 69.66 ±31.19 nmol/L, n = 8, fasted: 309.8 ±170.1 nmol/L, n = 8) and female mice (P <0.0001; control: 130.8 ±141.3 nmol/L, n =8, fasted: ±245.9 nmol/L, n = 8; Figure 1H). Furthermore, female fasted mice had greater CORT levels than male fasted mice (P<0.0001). Taken together, these results suggest that acute fasting decreased body weight and glucose concentration as well as increased ketones and CORT levels and the metabolic effect was more pronounced in female compared to male mice, regardless of weight loss.

### Sex differences in food approach behaviour and food consumption following fasting

To test if acute fasting altered food seeking differentially in male and female mice, we used a modified light dark box, whereby mice must enter the open light field to seek food (Figure 2A,B,E). Mice were weighed prior to being placed in the light dark box test. Male mice weighed more than female mice (sex effect: F(1,38) = 309.6, P < 0.0001; male control: 32 ± 2 g, n = 11; male fasted: 32 ± 0.4 g, n = 10; female control: 23 ±1 g, n = 11; female fasted: 22 ± 2 g, n = 10; Figure 2C). There was no main effect of fasting on distance travelled in the light zone (F(1,38) = 0.1770, P = 0.6763; Figure 2D,E). However, there was a main effect of sex on distance travelled in the light zone (F(1,38) = 9.853, P = 0.0033, sex x fasting interaction: F(1,38) = 0.0235, P = 0.9788; female control: 2398 ±410 cm, n = 11; female fasted: 2370 ±214 cm, n = 10, male control: 2082 ±363 cm, n = 11; male fasted: 2021 ±343 cm, n = 10, Figure 2D), suggesting that females have greater locomotor activity than males. Dark zone entries were not influenced by fasting (fasting effect: F(1,38) = 2.940, P = 0.0945, Figure 2E). However, there was a sex effect on dark zone entries (sex effect: F(1,38) = 7.063, P = 0.0114, fasting x sex interaction: : F(1,38) = 1.223, P = 0.2757, female control: 30 ± 10, n = 11; female fasted: 29 ± 7, n = 10, male control: 27 ± 4, n = 11; male fasted: 20 ± 7, n = 10). Although general activity in the light-dark box was not affect by fasting, behaviours within the food-zone were. There was a main effect of fasting on distance travelled in the food zone (F(1,38) = 11.15, P = 0.0019; Figure 2G). However, there was no sex difference on distance travelled in the food zone (sex effect: F(1,38) = 1.992, P = 0.1663, sex x fasting interaction: F(1,38) = 0.0017, P = 0.9967; male fasted: 155 ±59 cm, n = 10; female fasted: 180 ± 65cm, n = 10, male control: 93 ± 53 cm, n = 11; female control: 120 ± 58cm, n = 11; Figure 2G). There were also a greater number of food zone entries following fasting in all mice (fasting effect: F(1,38) = 13.64, P = 0.0007, male fasted: 37 ± 10, n = 10; female fasted: 43 ± 10, n = 10, male control: 26 ± 10, n = 11; female control: 31 ± 11, n = 11; Figure 2H). However, there was no effect of sex on food zone entries (sex effect: F(1,38) = 3.15, P = 0.08, sex x fasting interaction: main effect fasting: F(1,38) = 0.0495, P = 0.8250). The average duration of each food zone visit was greater following fasting (fasting effect: F(1,38) = 54.60, P <0.0001, male fasted: 2.3 ± 0.8 s, n = 10; female fasted: 1.5 ± 0.5 s, n = 10, control: 0.8 ± 0.3 s, n = 11; female control: 0.8 ± 0.3 s, n = 11; Figure 2I). There was a sex effect on the average duration in the food zone (sex effect: F(1,38) = 6.220, P = 0.0171, fasting x sex interaction: F(1,38) = 6.561, P = 0.0145). A Tukey’s multiple comparison test revealed a significant difference on the average duration in the food zone between male and female fasted mice (P = 0.0065), but not control mice (P >0.9999), suggesting that fasting influences the food seeking behaviour in female mice differently than male mice. During the 10 min test, fasted mice consumed more high-fat food during the test than control mice (fasting effect: F(1,36) = 108.8, P < 0.0001, male fasted: 0.26 ± 0.07 g, n = 10; female fasted: 0.17 ±0.09 g, n = 8, male control: 0.009 ±0.01 g, n = 11; female control: 0.028 ±0.04 g, n = 11; Figure 2J). Furthermore, there was fasting x sex interaction on food consumption (interaction: F(1,36) = 8.129, P = 0.0072). A Tukey’s multiple comparison test revealed significant differences in food consumption between control and fasted in both male (P <0.0001) and female mice (P<0.0001) as well as a difference between fasted male and female mice (P = 0.0161), but not in control mice (P = 0.8762), indicating that male mice consume more food during the test. This effect is consistent with greater food seeking behaviour observed in Figure 2I. During the 1 h period following the test, fasted mice consumed more kcal/body weight (g) of high fat diet than controls (fasting effect: F(1,38) = 23.90, P < 0.0001; male fasted: 0.15 ±0.02 kcal/g, n = 10; female fasted: 0.20 ±0.02 kcal/g, n = 10; male control: 0.12 ±0.03 kcal/g, n = 11; female control: 0.15 ±0.03 kcal/g, n = 11; Figure 2K). Similar to the behaviour observed in control mice during the 16 h isolation period, there was a sex effect in the kcal/g consumed (sex effect: F(1,38) = 24.79, P < 0.0001; interaction: F(1,38) = 0.5616, P = 0.4582). An a priori hypothesis that female mice consume more than males to restore energy balance was made based on control data (Figure 1C) and was tested with a Tukey’s multiple comparison test. There was a significant difference between male and female fasted mice (P = 0.0018; Figure 2K), suggesting that female fasted mice consumed more that male fasted mice after the test, consistent with their homeostatic baseline feeding pattern. Taken together, fasting increases food approach behaviours in male and female mice. However, male fasted mice spent more time in the food zone and consumed more food than female mice during the light dark box test.

### Effects of acute fasting on excitatory synaptic transmission

We next examined whether fasting altered basal synaptic properties at excitatory synapses onto VTA dopamine neurons. To assess the quantal release properties of excitatory inputs to VTA dopamine neurons, miniature excitatory post-synaptic currents (mEPSC) were recorded. There was a main effect of fasting (F(1,63) = 6.701, P = 0.0119) and sex (F(1,63) = 5.306, P = 0.0246) on mEPSC amplitude, but no fasting x sex interaction (F(1,63) = 1.309, P = 0.2568; male control: 15.6 ± 1.2 pA, N/n = 16/5, male fasted: 17.1 ±2.3 pA, N/n = 18/5; female control: 15.1 ±1.2 pA, N/n = 16/6, female fasted: 15.7 ±1.6 pA, N/n = 17/5; Figure 3C). A Tukey’s multiple comparison on the main effect of fasting indicated a significant difference in amplitude of male (P = 0.047), but not female mice (P = 0.741). Furthermore, there was a significant sex x fasting interaction on mEPSC frequency (interaction: F(1,63) = 5.150, P = 0.0267) as well as main effects of sex (F(1,63) = 9.281, P = 0.0034) and fasting (F(1,63) = 6.085, P = 0.014). Tukey’s multiple comparisons tests revealed fasting increased mEPSC frequency in male (P = 0.0068), but not female (P = 0.9999) mice compared to controls (male control: 0.5654 ±0.4270 Hz, N/n = 16/5; male fasted: 1.5 ± 1.3 Hz, N/n = 18/5; female control: 0.4 ± 0.4 Hz, N/n = 16/6; female fasted: 0.5 ± 0.3 Hz, N/n = 17/5, Figure 3D). These data indicate that fasting differentially affects mEPSC frequency onto VTA dopamine neurons of male and female mice.

An increase in the mEPSC frequency is commonly an indicator of presynaptic changes, which could be in the form of increased release probability (Pr) or increased number or function of the active release sites (n) (Simpson, 1971; Brock *et al*., 2020). To further examine glutamate release probability, the paired-pulse ratio (PPR) was calculated from evoked EPSCs from dopamine neurons of fasted and control mice. There were no significant group differences in PPR (sex x fasting interaction: F(1,40) = 0.8292, P = 0.3680, fasting effect: F(1,40) = 0.04698, P = 0.8295, sex effect: F(1,40) = 0.1747, P = 0.6782; male control: 0.7 ± 0.2, N/n = 12/3; male fasted: 0.7 ±0.1, N/n = 11/3; female control: 0.7 ±0.1, N/n = 10/4; female fasted: 0.7 ±0.2, N/n = 11/4; Figure 3E).

To test if fasting alters the size of the readily releasable pool and/or release probability, we used a short high frequency train stimulation of 40 pulses at 100 Hz (Thanawala & Regehr, 2013) (Figure 4A). Analysis of these evoked currents can reveal the relative size of the readily releasable pool (RRP(train)) and release probability (P(train)) (Figure 4B). RRP(train) was not changed by fasting (fasting effect: F(1,31) = 0.1113, P = 0.7409; male control: 1383 ±730, N/n = 9/4; male fasted: 1174 ±481, N/n = 11/3; female control: 1315 ±581, N/n = 8/2; female fasted: 1394 ±422, N/n = 7/2; Figure 4C). P(train) was also not changed following fasting (fasting effect: F(1,31) = 0.001955, P = 0.9650, male control: 0.3803 ±0.2468, N/n = 9/4; male fasted: 0.5 ±0.6, N/n = 11/3; female control: 0.3 ±0.2, N/n = 8/2; female fasted: 0.3 ±0.03, N/n = 7/2; Figure 4D). Furthermore, PPR calculated from these stimulations was not altered by fasting (fasting effect: F(1,31) = 0.08011, P = 0.7790; male control: 0.7791 ±0.3351, N/n = 9/4; male fasted: 0.8±0.3, N/n = 11/3; female control: 0.8 ± 0.2, N/n = 8/2; female fasted: 0.8 ±0.2, N/n = 7/2; Figure 4E). Taken together, these data indicate that fasting does not alter release probability or the size of the ready releasable pool at excitatory synapses onto dopamine neurons.

**Figure 4.**
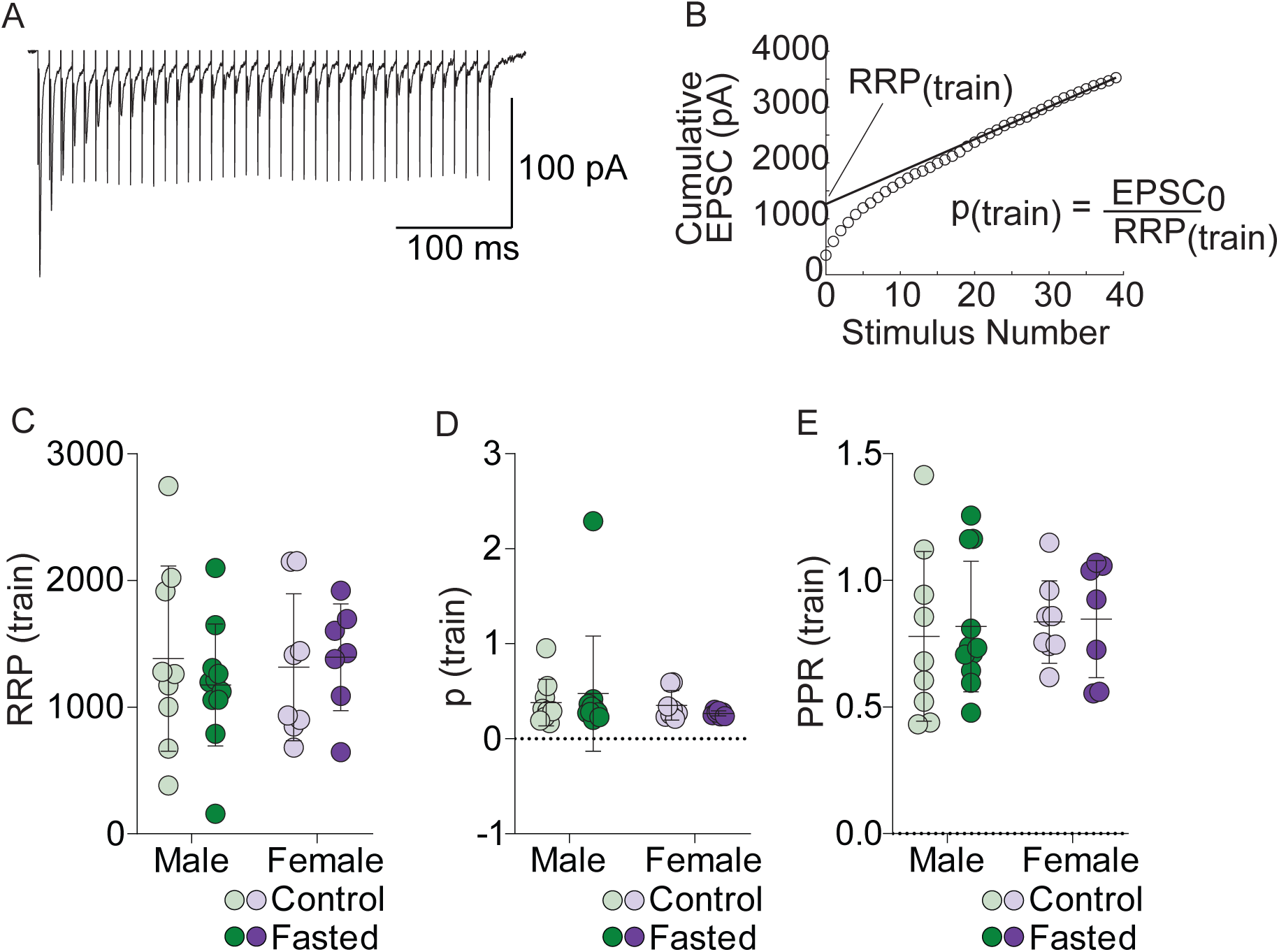
Effect of fasting on ready releasable pool and release probability from male and female mice. A) Representative example trace of 100 Hz train stimulation used for the readily releasable pool analysis. B) Example of the plotted cumulative amplitude and best fit line to determine the size of the readily releasable pool (RRP(train)) and the probability of release (P(train)). C) RRP determined from the y intercept of the fitted line of the cumulative amplitude. There was no effect of fasting or sex on the RRP. D) Release probability (P(train)) determined from the amplitude of the first evoked EPSC divided by RRP(train).There was no effect of fasting or sex on P(train). E) There was no effect of sex or fasting on PPR (train).

We next examined whether fasting altered synaptic efficacy of dopamine neurons through postsynaptic mechanisms. We first measured the AMPAR/NMDAR ratio, which reflects a relative change in the contribution of AMPAR, NMDAR or both. Fasting did not change the AMPAR/NMDAR ratio in male or female mice (fasting effect: F(1,32) = 0.01884, P = 0.8917; sex effect: F(1,32) = 1.268, P = 0.2686, male control: 0.6 ±0.3, N/n = 9/4; male fasted: 0.7 ±0.4, N/n = 10/5; female control: 0.8 ± 0.3, N/n = 8/4; female fasted: 0.7 ± 0.2,N/ n = 9/4; Figure 5A,B). To test if fasting altered the number or function of AMPARs that could contribute to increased mEPSC amplitude that now surpasses the threshold of event detection, we used Rubi-glutamate to uncage glutamate at excitatory synapses while recording AMPAR responses at -70 mV (Figure 5C). Responses to optical glutamate uncaging onto VTA dopamine neurons of male mice were not affected by fasting (t(34) = 0.1039, P = 0.9179, male fasted: 22.2 ±10.1 pA, N/n = 18/4, male control: 21.8 ±11.5 pA, N/n = 18/4; Figure 5D,E). Taken together, postsynaptic responses at glutamatergic synapses onto VTA dopamine neurons were not altered by fasting.

**Figure 5.**
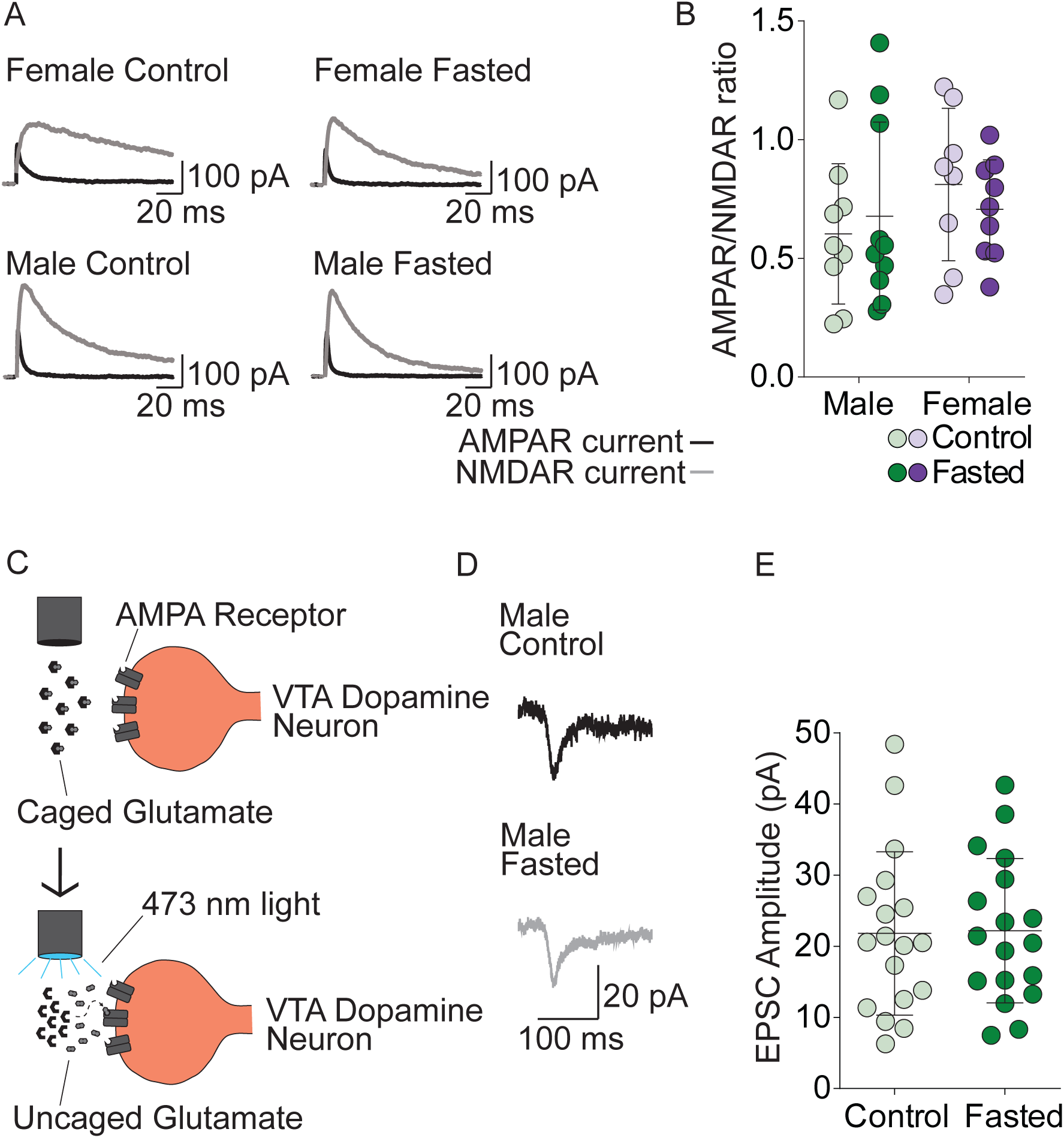
The effect of fasting on post-synaptic properties of excitatory synapses of male and female mice. A) Averaged representative traces of AMPAR and NMDAR EPSCs. B) The AMPAR/NMDAR ratio of fasted and control male and female mice. There was no effect of fasting or sex on the AMPAR/NMDA ratio. C) Schematic illustrating the optically-induced (470 nm) uncaging of glutamate using RuBi-Glutamate. D) Averaged representative traces of glutamate currents recorded at -70 mV. E) EPSC amplitude in response to photo-uncaging of Rubi-glutamate in control or fasted male mice. There was no difference of EPSCs onto dopamine neurons from control or fasted male mice.

### Effect of fasting on depolarization-induced suppression of excitation

Acute fasting could be considered a physiological stressor to mice, as was reflected in increased CORT levels in female mice. In response to glucocorticoids, endogenous cannabinoids (endocannabinoids) induce a rapid suppression of synaptic transmission in the hypothalamus and the basolateral amygdala (Di *et al*., 2003, 2016). Endocannabinoids act as retrograde transmitters that are synthesized and released on demand and act presynaptically at cannabinoid 1 (CB1) receptors to suppress neurotransmitter release (Wilson & Nicoll, 2002) (Figure 6A). To test if fasting differentially altered endocannabinoid signalling in males and females, we examined depolarization-induced suppression of excitation (DSE), a transient form of plasticity mediated by endocannabinoids. We recorded evoked EPSCs before and after a 10s depolarization. DSE was abolished by the CB1 receptor antagonist, AM251, in both males (drug effect: F(1,52) = 8.492, P = 0.0052, male control with AM251: 96.8 ± 19.3 % baseline, N/n = 5/2; male fasted with AM251: 108.3 ±7.5 % baseline, N/n = 4/2; Figure 6C,D) and females (drug effect: F(1,58) = 40.27, P < 0.0001, female control with AM251: 94.6 ±11.7 % baseline, N/n = 8/4; female fasted with AM251: 113.2 ± 9.9 % baseline, N/n = 10/4; Figure 6F,G), confirming that DSE is endocannabinoid mediated. There was no effect of fasting on DSE in the VTA of male mice (fasting effect: F(1,52) = 0.7370, P = 0.3946, male control: 76.6 ± 23. 9 % baseline, N/n = 25/10; male fasted: 79.56 ±23. 9 % baseline, N/n = 22/8; Figure 6B,D). However, there was a significant fasting x drug interaction on DSE in female mice (interaction: F(1,58) = 12.22, P = 0.0009, female control: 79.8 ± 22.7 % baseline, N/n = 24/13; female fasted: 61.9 ± 18.1 % baseline, N/n = 20/8; Figure 6E,G). A Tukey’s multiple comparisons test indicated a significant difference in DSE between fasted and control female mice in the absence (P = 0.0119), but not in the presence of AM251 (P = 0.1617). When male and female mice were compared together, there was a significant sex x fasting interaction on DSE (interaction: F(1,87) = 4.890, P = 0.0296). A Tukey’s multiple comparisons test indicated a significant difference between female fasted and control mice (P = 0.0482), but not male fasted and control mice (P = 0.9683, Figure 6I). Taken together, fasting increases DSE in female, but not male mice.

**Figure 6.**
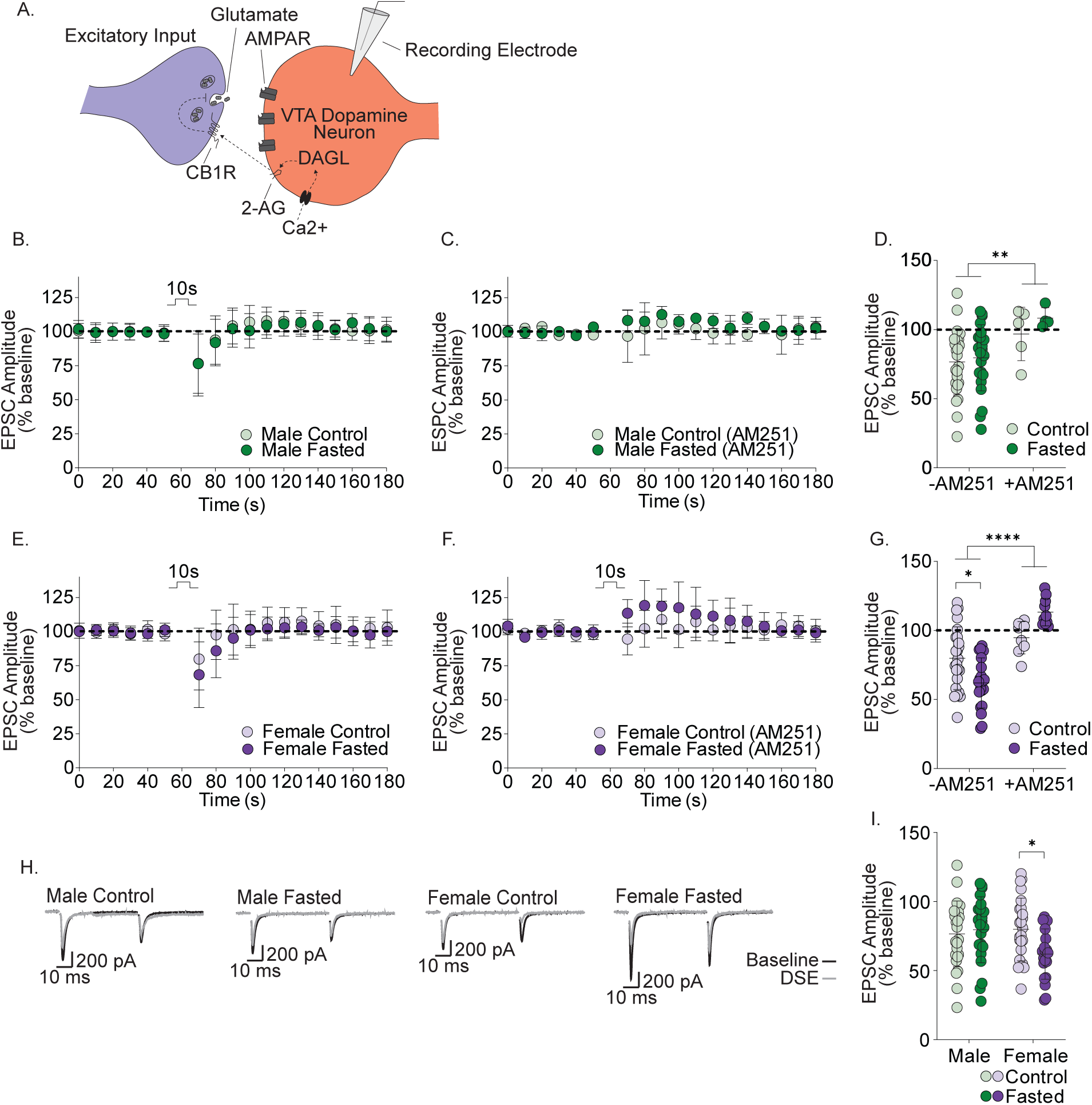
Sex differences in DSE of dopamine neurons from fasted and control male and female mice. A) Schematic illustrating the endocannabinoid mediated depolarization induced suppression of excitation (DSE). B) Time course of evoked EPSCs before and after the DSE protocol in dopamine neurons of male control and fasted mice. C) Time course of evoked EPSCs in the presence of the CB1 receptor antagonist AM251 (2 µM) before and after the DSE protocol in dopamine neurons of male control and fasted mice. D) Averaged responses during DSE of dopamine neurons from male control or fasted mice in the presence or absence of AM251. AM251 inhibits DSE in male control or fasted mice. E) Time course of evoked EPSCs before and after the DSE protocol in dopamine neurons of female control and fasted mice. F) Time course of evoked EPSCs in the presence of the CB1 receptor antagonist AM251 (2 µM) before and after the DSE protocol in dopamine neurons of female control and fasted mice. G) Averaged responses during DSE of dopamine neurons from female control or fasted mice in the presence or absence of AM251. DSE was greater in fasted female mice compared to controls. This effect was blocked by AM251. H) Averaged representative traces of EPSCs during baseline and DSE of dopamine neurons from male and female fasted and control mice. I) Averaged responses during DSE of dopamine neurons from female control or fasted mice.

To determine if the increased DSE in female mice is due to an increase in the CB1 receptor-expression or a change in receptor efficacy, we applied a maximal concentration of CB1 receptor agonist, WIN 55,212 (10 µM) to VTA slices. If there was an increase in receptor expression or receptor efficacy, the maximal response to the agonist should be greater. Although WIN 55,212 inhibited EPSCs in the VTA of female mice (drug effect: F(1,6) = 66.71, P = 0.0002), this effect was not altered by fasting (fasting effect: F(1,6) = 0.1244, P = 0.7364, control: 79.7 ± 4.3 % baseline, N/n = 4/4; fasted: 77.9 ± 9.5 % baseline, N/n = 4/3; Figure 7A-C). Consistent with a presynaptic locus, WIN 55,212 increased PPR (drug effect: F(1,6) = 10.90, P = 0.0164; control PPR baseline: 0.75 ±0.2, N/n = 4/4; control PPR WIN 55,212: 0.82 ±0.2, fasted PPR baseline: 0.79 ±0.1, N/n = 4/4, fasted PPR WIN 55,212: 0.87 ±0.05, N/n = 4/4; Figure 7D). Taken together, increase in DSE observed in female mice after fasting was not due to a fasting-induced change in receptor number and/or efficacy.

**Figure 7.**
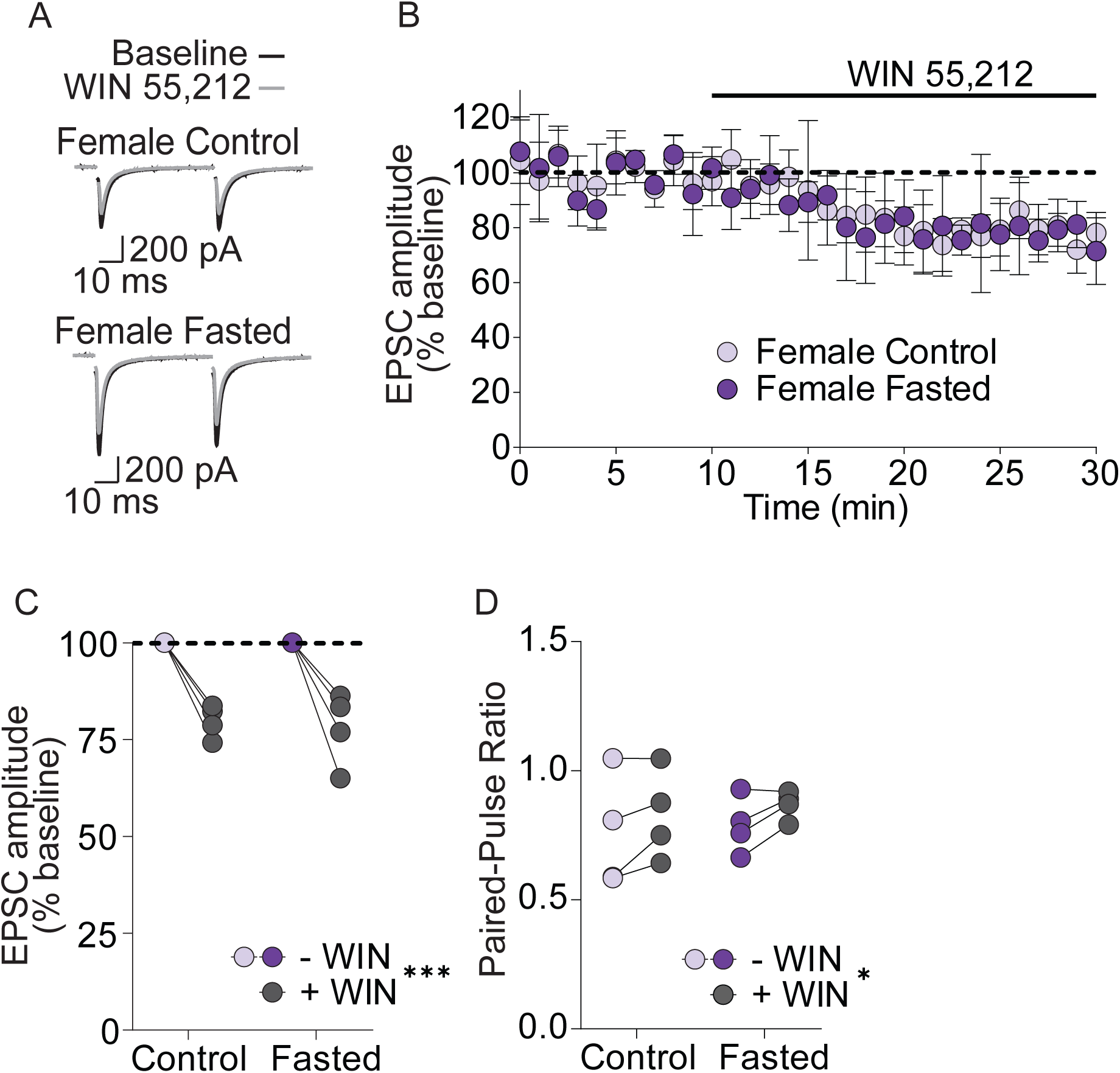
The CB1 receptor agonist, WIN 55,212, supresses EPSCs but responses are not altered by fasting. A) Averaged example traces of EPSCs of dopamine neurons from female during baseline and after the application of WIN 55,212. B) Time course of evoked EPSCs before, during and after WIN 55,212 (10 µM) application to dopamine neurons of female control and fasted mice. C) Averaged responses of EPSCs before and after WIN 55,212 application to dopamine neurons from female control or fasted mice. WIN 55,212 decreased EPSC amplitude in fasted and control mice. D) PPR before and after WIN 55,212 application to dopamine neurons of control and fasted female mice. WIN 55,212 increased PPR in fasted and control mice.

### Effects of fasting at inhibitory synapses in the VTA

We next examined the effect of fasting on GABAergic synapses onto VTA dopamine neurons of male and female mice. To assess the quantal release properties of inhibitory synapses, we recorded miniature inhibitory post-synaptic currents (mIPSC) in control and fasted mice. There was no effect of fasting on mIPSC amplitude (fasting effect: F(1,62) = 1.137, P = 0.2905) nor was there an effect of sex on mIPSC amplitude (sex effect: F(1,62) = 0.0615, P = 0.805, male control: 21.9 ±1.9 pA, N/n = 17/5; female control: 21.6 ±1.7 pA, N/n = 17/6; male fasted: 22.2 ±1.8 pA, N/n = 17/5; female fasted: 22.3 ±1.9 pA, N/n = 15/5; Figure 8A,B). Furthermore, there was no effect of fasting on mIPSC frequency (fasting effect: F(1,62) = 0.8943, P = 0.3480). There was a potential trending effect of sex on mIPSC frequency (sex effect: F(1,62) = 3.463, P = 0.0675, male control: 1.6 ±1.3 Hz, N/n = 17/5; female control: 1.1 ±0.5 Hz, N/n = 17/6; male fasted: 1.9 ±1.6 Hz, N/n = 17/5; female fasted: 1.4 ±1.1 Hz, N/n = 16/5; Figure 8A,C).

**Figure 8.**
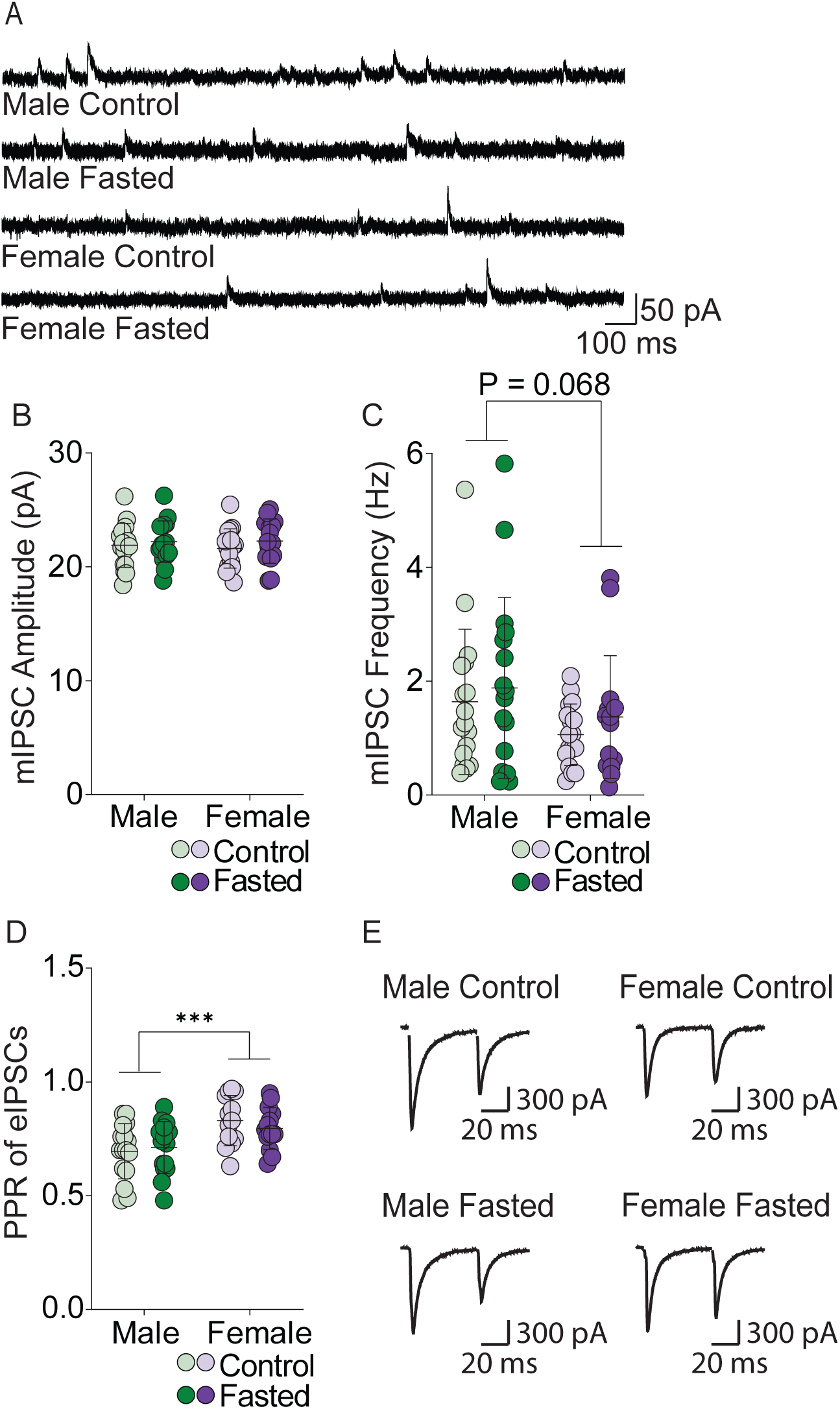
Quantal release properties at inhibitory synapses onto dopamine neurons of male or female mice do not change following fasting. A) Representative examples of mIPSC recordings from male and female fasted and control mice. B) mIPSC amplitude onto dopamine neurons of male and female fasted and control mice. There was no effect of fasting or sex on mIPSC amplitude. C) mIPSC frequency onto dopamine neurons of male and female fasted and control mice. D) Paired-pulse ratio of evoked IPSCs was not different between fasted or control male and female mice. Female control and fasted mice had a higher PPR compared to male mice. E) Representative example recordings of the paired-pulse evoked IPSC currents.

Because we observed a trend in sex difference on mIPSC frequency, we further examined the PPR of evoked IPSCs, a measure associated with release probability. There was no fasting effect on PPR (fasting effect: F(1,54) = 0.08796, P = 0.7679; Figure 8D,E). However, there was a significant effect of sex on PPR at inhibitory synapses (sex effect: F(1,54) = 13.98, P = 0.0004; female control: 0.8 ±0.1, N/n = 13/4; female fasted: 0.8 ±0.09, N/n = 14/4, male control: 0.7 ±0.1, N/n = 16/6; male fasted: : 0.7 ±0.1, N/n = 15/5, Figure 8D,E). Taken together, fasting does not alter release probability at inhibitory synapses in male or female mice. However, female mice may have lower release probability at inhibitory synapses compared to male mice.

### Effect of fasting on depolarization-induced suppression of inhibition

Depolarization-induced suppression of inhibition (DSI) was used to assess endocannabinoid-mediated short-term presynaptic depression at inhibitory synapses. There was a main effect of fasting on DSI of VTA dopamine neurons of male (fasting effect: F(1,42) = 4.314, P = 0.044, male control: 73.8 ±12.8% baseline, N/n = 16/6; male fasted: 61.7 ±18.0 % baseline, N/n = 15/4; Figure 9A,C), but not female mice (fasting effect: F(1,37) = 3.218, P = 0.08, female control: 59.8 ±13.6 % baseline, N/n = 13/4; female fasted: 72.5 ±17.0 % baseline, N/n = 14/4; Figure 9D,F). DSI was inhibited by AM251 in both males (drug effect: F(1,42) = 10.97, P = 0.0019, male control with AM251: 84.5 ±8.1 % baseline, N/n = 7/2; male fasted with AM251: 79.02 ±5.143 % baseline, N/n = 8/3; Figure 9B,C) and females (drug effect: F(1,37) = 29.69, P < 0.0001, female control with AM251: 88.6 ±7.5 % baseline, N/n = 6/2; female fasted with AM251: 91.6 ±5.7 % baseline, N/n = 8/3; Figure 9E,F). When sexes were compared directly, there was significant sex x fasting interaction on DSI (interaction: F(1,54) = 9.245, P = 0.0036; Figure 9H). However, there were no significant group differences (sex effect: F(1,54) = 0.1525, P = 0.6977). Taken together, fasting did not induce robust effects on DSI in male or female mice.

**Figure 9.**
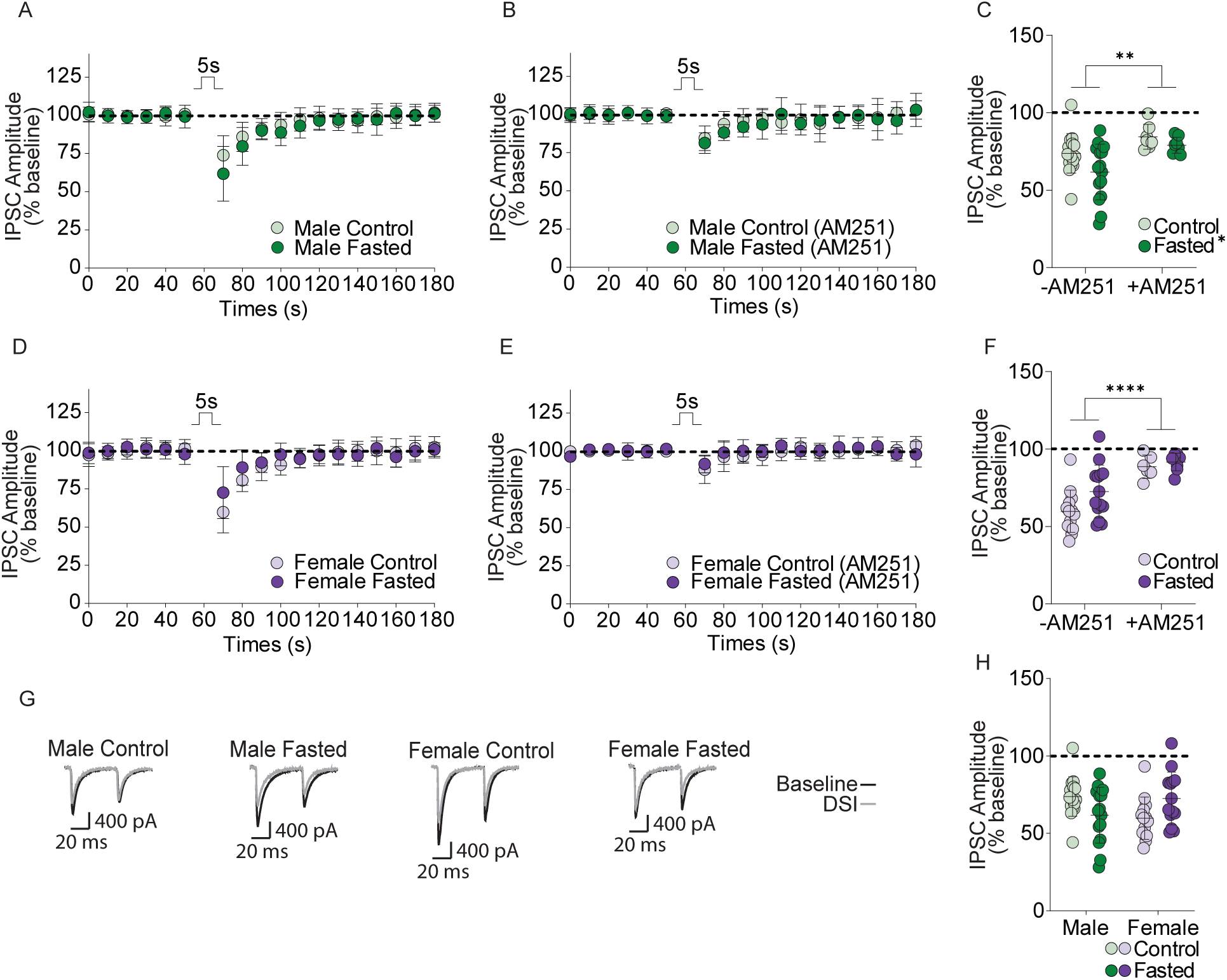
Fasting does not alter DSI at inhibitory synapses onto dopamine neurons of male or female mice. A) Time course of evoked IPSCs before and after the DSI protocol in dopamine neurons of male control and fasted mice. B) Time course of evoked IPSCs in the presence of the CB1 receptor antagonist AM251 (2 µM) before and after the DSI protocol in dopamine neurons of male control and fasted mice. C) Averaged responses during DSI of dopamine neurons from male control or fasted mice in the presence or absence of AM251. AM251 inhibited DSI in control and fasted male mice. D) Time course of evoked IPSCs before and after the DSI protocol in dopamine neurons of female control and fasted mice. E) Time course of evoked IPSCs in the presence of the CB1 receptor antagonist AM251 (2 µM) before and after the DSI protocol in dopamine neurons of female control and fasted mice. F) Averaged responses during DSI of dopamine neurons from female control or fasted mice in the presence or absence of AM251. AM251 inhibited DSI in control and fasted female mice. G) Averaged representative traces of IPSCs during baseline and DSI of dopamine neurons from male and female fasted and control mice. H) Averaged responses during DSI of dopamine neurons from female control or fasted mice. There is no effect of sex or fasting on DSI.

## Discussion

We observed that an acute overnight 16 h fast induced strong metabolic and behavioural changes that affected females differently from males. Females had lower blood glucose, higher ketones and higher CORT levels than male mice. Furthermore, female mice had increased locomotor activity in general, but not in the food zone and consumed less food during the test than male mice. While fasting increased food-seeking in both male and female mice, synaptic transmission at inhibitory synapses in the VTA was preserved with only subtle sex differences observed at excitatory synapses. Dopamine neurons from male mice had an increase in mEPSC amplitude and frequency after fasting, whereas female mice had increased endocannabinoid-mediated short-term plasticity at excitatory synapses onto dopamine neurons. Given that many procedures involve acute fasting (i.e. measurement of fasted glucose, initiation of operant conditioning), it is important to understand how this may influence synaptic processes underlying reward learning.

There is a significant variability in the literature associated with physiological effects of acute fasting. This is, in part, due to variability of the length of the fast, the level of food restriction (i.e. total vs. partial), and the timing within the circadian rhythm (Jensen *et al*., 2013). The fasting protocol used here is consistent with fasting models used in other studies and yielded consistent physiological effects associated with an acute fast (Jensen *et al*., 2013). This procedure allowed for access to water and fasting over the mouse’s active (dark) phase. Consistent with similar fasting protocols (Jensen *et al*., 2013), we observed a 10% reduction in body weight. Weight loss associated with this duration of fast is due both to a reduction in body weight (60%) and gastric emptying (40%) (Prior *et al*., 2012). In addition, decreased blood glucose, likely due to decreased glucose absorbed from the gastrointestinal tract during the fast, as well as increased ketones is consistent with previous work (Jensen *et al*., 2013). Moreover, our results replicate previous work using an overnight fast demonstrating increased CORT in fasted females compared to males (Champy *et al*., 2004). Thus, a 16 h overnight fast used in our model induces reliable physiological effects consistent with other studies.

Acute fasting increased food approach behaviours predominantly in male mice. Notably, increased food approach behaviours are associated with increased mEPSC frequency and excitatory synapse formation onto VTA dopamine neurons (Liu *et al*., 2016). Female mice had increased overall locomotor activity and dark zone entries compared to male mice in addition to higher fasting-induced CORT levels on par with CORT induction from other types of stressors (Bowers *et al*., 2008). Notably, while males are more likely to engage with the food during the light dark box test, female fasted mice consumed more food than males in the 1h period after the test. This effect may be a function of higher CORT levels in female mice, given that CORT can drive food intake (Castonguay *et al*., 1986). Therefore, it is feasible that fasting is a stronger physiological stressor in female than male mice and this stress may contribute to reduced food consumption during the task compared to male mice as well as increased homeostatic food intake after the task.

We observed a fasting-induced increase in mEPSC frequency and amplitude in male, but not female mice. Increased mEPSC frequency is typically associated with increased glutamate release (Simpson, 1971; Holz & Fisher, 1999; Brock *et al*., 2020). However, we did not observe changes in release probability or the size of the readily releasable pool. Furthermore, it is unlikely that the fasting-induced increase in mEPSC frequency is due to an increase in amplitude of AMPAR response that now passes the threshold for mEPSC detection as we did not observe a change in AMPAR/NMDAR ratio or responses to Rubi-glutamate. Thus, fasting does not induce a postsynaptic change in number or function of AMPARs or a post-synaptic unsilencing of glutamatergic synapses in the VTA. An alternative explanation for increased mEPSC frequency could be due to an increase in the number of release sites, which could be measured with immuno-electron microscopy of excitatory synapses. Notably, an increase in ketones during fasting is associated with increased brain derived neurotrophic factor (BDNF) in the cortex and hippocampus (Marosi *et al*., 2016; Sleiman *et al*., 2016; Mattson *et al*., 2018), which can induce formation of new synaptic release sites in the VTA (Pu *et al*., 2006). Exposure to sweetened high fat food increased excitatory synapse number onto VTA dopamine neurons, measured with electrophysiology and immuno-electron microscopy. This effect was associated with increased food-seeking behaviour in a light-dark box test (Liu *et al*., 2016). Decreasing excitatory synaptic strength in the VTA reverses the palatable-food increase in food approach behaviours (Liu *et al*., 2016). Thus, increased excitatory synapse number onto VTA dopamine neurons of male mice may underlie enhanced food seeking, regardless of whether this is driven by energy density or by food-restriction.

Fasting increased endocannabinoid-mediated DSE at excitatory synapses onto dopamine neurons of female, but not male mice. Others have demonstrated that a 24-h fast induced a loss of endocannabinoid-mediated LTD at GABAergic synapses in the dorsomedial hypothalamus, an effect mediated by glucocorticoids (Crosby *et al*., 2011). Furthermore, increased CORT is associated with increased endocannabinoids in the amygdala or hypothalamus (Di *et al*., 2003, 2016). A fasting-induced facilitation of DSE was not associated with a change in basal PPR or ready releasable pool in female mice indicating that fasting does not induce a lasting endocannabinoid tone. Furthermore, we did not observe altered CB1 receptor efficacy with fasting of female mice. Endocannabinoid synthesis and release is initiated on demand by an influx in calcium and is influenced by expression of enzymes including d-acyl glycerol lipase (DAGL) and n-arachidonylphosphatidylethanolamine phospholipase D (NAPE-PLD). Future experiments could test if fasting alters expression or function of DAGL or NAPE-PLD differentially in male or female mice. Taken together, fasting-induced changes in excitatory inputs of dopamine neurons of female mice may be expressed under more depolarized conditions allowing for calcium influx and endocannabinoid synthesis.

Fasting did not induce changes at inhibitory synapses onto VTA dopamine neurons of male or female mice. As both DSE and DSI are mediated by 2-AG in the VTA, it is interesting that only DSE is affected by fasting in female mice. This finding is not without precedence, as insulin induces endocannabinoid-mediated LTD of excitatory, but not inhibitory, synapses in the VTA (Labouèbe *et al*., 2013). This difference could be due to spatial segregation of glutamatergic and GABAergic synapses within the VTA, whereby endocannabinoids produced near glutamatergic synapses many not reach GABAergic synapses. Notably, there is strong segregation of GABAergic input to distal dendrites of dopamine neurons that extend into the substantia nigra reticulata whereas there is a higher proportion of glutamatergic synapses near proximal dendrites and perikaryon in the substantia nigra pars compacta (Henny *et al*., 2012). Although fasting had no effect on the inhibitory synapses, we did observe that the probability of release at inhibitory synapses onto VTA neurons was lower in female compared to male mice, regardless of fasting, suggesting that there may be basal differences in release properties.

In summary, acute fasting affects metabolic parameters, food-seeking, and synaptic properties in the VTA in a sex-dependent manner. While a 16 h overnight fast increased food-seeking behaviour overall supporting the efficacy of the fasting procedure, this increase was significantly greater in male mice. Furthermore, male mice had increased mEPSC frequency onto VTA dopamine neurons, an effect likely due to increased number of release sites. Female mice on the other hand showed no changes in basal synaptic properties, but rather a facilitation in endocannabinoid-mediated short-term depression at excitatory synapses onto dopamine neurons. Inhibitory synapses onto dopamine neurons of male or female mice were not changed by fasting. Thus, a 16h fast leads to sexually dimorphic synaptic changes that parallel food-seeking behaviour. These results highlight the sensitivity and importance of VTA dopamine neurons in responding to changes in motivational states. Furthermore, fasting-induced plasticity in the VTA may underlie enhanced food-seeking during food availability and thus, may be a reason why weight loss is rarely maintained. Importantly, major depressive disorder and anxiety are twice as frequently diagnosed in women compared to men, while anorexia nervosa is 13 times more frequent in women (McCarthy *et al*., 2012). Therefore, understanding sex differences in state dependent changes in synaptic function of VTA dopamine neurons may highlight mechanisms underlying why there is gender and sex bias in these neuropsychiatric illnesses.

## Acknowledgments

This work is supported by a Tier 1 Canada Research Chair (950-232211) and Canada Institutes for Health Research Foundation Grant (CIHR FDN 148473 SLB). NG was supported by a Harley Hotchkiss Graduate Scholarship. The authors wish to acknowledge the behavioural core facility at the Hotchkiss Brain Institute for use of their light-dark box.

## Author Contributions

NG performed electrophysiological and behavioural experiments. NG and SLB wrote first drafts of the manuscript. NG and SLB revised the manuscript.

## Data Availability

Data is available upon request to the corresponding author.

## References

Borgland SL, Taha SA, Sarti F, Fields HL & Bonci A (2006). Orexin a in the VTA is critical for the induction of synaptic plasticity and behavioral sensitization to cocaine. Neuron 49, 589–601.

Bowers SL, Bilbo SD, Dhabhar FS & Nelson RJ (2008). Stressor-Specific Alterations in Corticosterone and Immune Responses in Mice. Brain, Behav Immun 22, 105–113.

Branch SY, Goertz RB, Sharpe AL, Pierce J, Roy S, Ko D, Paladini CA & Beckstead MJ (2013). Food restriction increases glutamate receptor-mediated burst firing of dopamine neurons. J Neurosci 33, 13861–13872.

Brock JA, Thomazeau A, Watanabe A, Li SSY & Sjöström PJ (2020). A Practical Guide to Using CV Analysis for Determining the Locus of Synaptic Plasticity. Front Synaptic Neurosci 12, 1–16.

Cadoni C, Solinas M, Valentini V & Di Chiara G (2003). Selective psychostimulant sensitization by food restriction: Differential changes in accumbens shell and core dopamine. Eur J Neurosci 18, 2326–2334.

Carr KD (2002). Augmentation of drug reward by chronic food restriction: Behavioral evidence and underlying mechanisms. Physiol Behav 76, 353–364.

Castonguay TW, Dallman MF, Stern JS (1986). Some metabolic and behavioural effects of adrenalectomy on obese Zucker rats. Am J Physiol Reg Int Comp Physiol 251, R923–R933.

Champy MF, Selloum M, Piard L, Zeitler V, Caradec C, Chambon P & Auwerx J (2004). Mouse functional genomics requires standardization of mouse handling and housing conditions. Mamm Genome 15, 768–783.

Crosby KM, Inoue W, Pittman QJ & Bains JS (2011). Endocannabinoids gate state-dependent plasticity of synaptic inhibition in feeding circuits. Neuron 71, 529–541.

Di S, Itoga CA, Fisher MO, Solomonow J, Roltsch EA, Gilpin NW & Tasker JG (2016). Acute stress suppresses synaptic inhibition and increases anxiety via endocannabinoid release in the basolateral amygdala. J Neurosci 36, 8461–8470.

Di S, Malcher-Lopes R, Halmos KC & Tasker JG (2003). Nongenomic glucocorticoid inhibition via endocannabinoid release in the hypothalamus: A fast feedback mechanism. J Neurosci 23, 4850–4857.

Fino E, Araya R, Peterka DS, Salierno M, Etchenique R & Yuste R (2009). RuBi-Glutamate: Two-photon and visible-light photoactivation of neurons and dendritic spines. Front Neural Circuits 3, 1–9.

La Fleur SE, Vanderschuren LJMJ, Luijendijk MC, Kloeze BM, Tiesjema B & Adan RAH (2007). A reciprocal interaction between food-motivated behavior and diet-induced obesity. Int J Obes 31, 1286–1294.

Freire T, Senior AM, Perks R, Pulpitel T, Clark X, Brandon AE, Wahl D, Hatchwell L, Le Couteur DG, Cooney GJ, Larance M, Simpson SJ & Solon-Biet SM (2020). Sex-specific metabolic responses to 6 hours of fasting during the active phase in young mice. J Physiol 598, 2081–2092.

Fulton S (2010). Appetite and reward. Front Neuroendocrinol 31, 85–103.

Grundy D (2015). Principles and standards for reporting animal experiments in The Journal of Physiology and Experimental Physiology. J Physiol 593, 2547–2549.

Heffner TG, Hartman JA & Seiden LS (1980). Feeding increases dopamine metabolism in the rat brain. Science (80-) 208, 1168–1170.

Henny P, Brown MTC, Northrop A, Faunes M, Ungless MA, Magill PJ & Bolam JP (2012). Structural correlates of heterogeneous in vivo activity of midbrain dopaminergic neurons. Nat Neurosci 15, 613–619.

Holz RW & Fisher SK (1999). Synaptic Transmission. In Basic Neurochemistry: Molecular, Cellular and Medical Aspects, 6th editio., ed. Siegel G, Agranoff B & Albers R. Lippincott-Raven, Philadelphia. Available at: https://www.ncbi.nlm.nih.gov/books/NBK27911/.

Jensen TL, Kiersgaard MK, Sørensen DB & Mikkelsen LF (2013). Fasting of mice: A review. Lab Anim 47, 225–240.

Jewett DC, Cleary J, Levine AS, Schaal DW & Thompson T (1995). Effects of neuropeptide Y, insulin, 2-deoxyglucose, and food deprivation on food-motivated behavior. Psychopharmacology (Berl) 120, 267–271.

Johnstone AM, Faber P, Gibney ER, Elia M, Horgan G, Golden BE & Stubbs RJ (2002). Effect of an acute fast on energy compensation and feeding behaviour in lean men and women. Int J Obes 26, 1623–1628.

Julia C, Péneau S, Andreeva VA, Méjean C, Fezeu L, Galan P & Hercberg S (2014). Weight-loss strategies used by the general population: How are they perceived? PLoS One 9, 1–8.

Labouèbe G, Liu S, Dias C, Zou H, Wong JCY, Karunakaran S, Clee SM, Phillips AG, Boutrel B & Borgland SL (2013). Insulin induces long-term depression of ventral tegmental area dopamine neurons via endocannabinoids. Nat Neurosci 16, 300–308.

Liu S, Globa AK, Mills F, Naef L, Qiao M, Bamji SX & Borgland SL (2016). Consumption of palatable food primes food approach behavior by rapidly increasing synaptic density in the VTA. Proc Natl Acad Sci U S A 113, 2520–2525.

Marosi K, Kim SW, Moehl K, Scheilbey-Knudsen M, Cheng A, Cutler R, Camandola S & Mattson MP (2016). 3-Hydroxybutyrate Regulates Energy Metabolism and Induces BDNF Expression in Cerebral Cortical Neurons. J Neurochem 139, 769–781.

Matikainen-Ankney BA, Ali MA, Miyazaki NL, Fry SA, Licholai JA & Kravitz A V. (2020). Weight Loss After Obesity is Associated with Increased Food Motivation and Faster Weight Regain in Mice. Obesity 28, 851–856.

Mattson MP, Moehl K, Ghena N, Schmaedick M & Cheng A (2018). Intermittent metabolic switching, neuroplasticity and brain health. Nat Rev Neurosci 19, 63–80.

McCarthy MM, Arnold AP, Ball GF, Blaustein JD & de Vries GJ (2012). Sex differences in the brain: The not so inconvenient truth. J Neurosci 32, 2241–2247.

Melis M, De Felice M, Lecca S, Fattore L & Pistis M (2013). Sex-specific tonic 2-arachidonoylglycerol signaling at inhibitory inputs onto dopamine neurons of Lister hooded rats. Front Integr Neurosci 7, 1–13.

Melis M, Pistis M, Perra S, Muntoni AL, Pillolla G & Gessa GL (2004). Endocannabinoids Mediate Presynaptic Inhibition of Glutamatergic Transmission in Rat Ventral Tegmental Area Dopamine Neurons through Activation of CB1 Receptors. J Neurosci 24, 53–62.

Paterson TA, Brot MD, Zavosh A, Schenk JO, Szot P & Figlewicz DP (1998). Food Deprivation Decreased mRNA and Activity of the Rat Dopamine Transporter. Neuro-endorcinology 68, 11–20.

Prior H, Ewart L, Bright J & Valentin JP (2012). Refinement of the charcoal meal study by reduction of the fasting period. ATLA Altern to Lab Anim 40, 99–107.

Pu L, Liu QS & Poo MM (2006). BDNF-dependent synaptic sensitization in midbrain dopamine neurons after cocaine withdrawal. Nat Neurosci 9, 605–607.

Reilly S (1999). Reinforcement value of gustatory stimuli determined by progressive ratio performance. Pharmacol Biochem Behav 63, 301–311.

Roseberry AG (2015). Acute fasting increases somatodendritic dopamine release in the ventral Tegmental area. J Neurophysiol 114, 1072–1082.

Salamone JD & Correa M (2012). The Mysterious Motivational Functions of Mesolimbic Dopamine. Neuron 76, 470–485.

Sharma S, Fernandes MF & Fulton S (2013). Adaptations in brain reward circuitry underlie palatable food cravings and anxiety induced by high-fat diet withdrawal. Int J Obes 37, 1183–1191.

Simpson JA (1971). Electrophysiological Analysis of Synaptic Transmission.

Sleiman SF, Henry J, Al-Haddad R, El Hayek L, Haidar EA, Stringer T, Ulja D, Karuppagounder SS, Holson EB, Ratan RR, Ninan I & Chao M V. (2016). Exercise promotes the expression of brain derived neurotrophic factor (BDNF) through the action of the ketone body β-hydroxybutyrate. Elife 5, 1–21.

Stuber GD, Klanker M, De Ridder B, Bowers MS, Joosten RN, Feenstra MG & Bonci A (2008). Reward-predictive cues enhance excitatory synaptic strength onto midbrain dopamine neurons. Science (80-) 321, 1690–1692.

Thanawala MS & Regehr WG (2013). Presynaptic calcium influx controls neurotransmitter release in part by regulating the effective size of the readily releasable pool. J Neurosci 33, 4625–4633.

Thanawala MS & Regehr WG (2016). Determining synaptic parameters using high-frequency activation. J Neurosci Methods 264, 136–152.

Wilson C, Nomikos GG, Collu M & Fibiger HC (1995). Dopaminergic correlates of motivated behavior: Importance of drive. J Neurosci 15, 5169–5178.

Wilson RI & Nicoll RA (2002). Neuroscience: Endocannabinoid signaling in the brain. Science (80-) 296, 678–682.

Wing RR, Lang W, Wadden TA, Safford M, Knowler WC, Bertoni AG, Hill JO, Brancati FL, Peters A & Wagenknecht L (2011). Benefits of modest weight loss in improving cardiovascular risk factors in overweight and obese individuals with type 2 diabetes. Diabetes Care 34, 1481–1486.

